# Synchrotron FTIR and Raman spectroscopy 1 ectroscopy provide unique spectral fingerprints for *Arabidopsis* floral stem vascular tissues

**DOI:** 10.1101/343343

**Authors:** S Dinant, N Wolff, F De Marco, F Vilaine, L Gissot, E Aubry, C Sandt, C Bellini, R Le Hir

## Abstract

Cell walls are highly complex structures that are modified during plant growth and development. For example, the development of phloem and xylem vascular cells, which participate in the transport of sugars and water as well as support, can be influenced by cell-specific cell wall composition. Here, we used synchrotron radiation-based infrared (SR-FTIR) and Raman spectroscopy to analyze the cell wall composition of wild-type and double mutant *sweet11-1sweet12-1*, which impairs sugar transport, Arabidopsis floral stem vascular tissue. The SR-FTIR spectra showed that in addition to modified xylem cell wall composition, phloem cell walls in the double mutant line were characterized by modified hemicellulose composition. Moreover, combining Raman spectroscopy with a Classification and Regression Tree (CART) method identified combinations of Raman shifts that could distinguish xylem vessels and fibers. Additionally, the disruption of *SWEET11* and *SWEET12* genes impacts xylem cell wall composition in a cell-specific manner, with changes in hemicelluloses and cellulose observed at the xylem vessel interface. These results suggest that the facilitated transport of sugars by transporters that exist between vascular parenchyma cells and conducting cells is important to ensuring correct phloem and xylem cell wall composition.

**Highlight:** Combining vibrational spectroscopy techniques and multivariate analysis shows that the disruption of *SWEET* genes impacts phloem cell wall composition and that the effect on xylem cell wall composition is cell-specific.

## Introduction

The presence of a polysaccharide-rich frame is an important feature of plant cells. The primary cell wall, composed mainly of insoluble (cellulose and hemicelluloses) and soluble polysaccharides (pectins), is deposited when plant cells are growing. Once the cells stop growing, the primary cell wall is reinforced by a secondary cell wall (SCW), which is composed mainly of cellulose, hemicelluloses and lignin. To ensure their specialized function in structural support and water transportation, the xylem vessels and fibers have an even thicker SCW. Secondary cell wall production in plant cells is of interest to humans because it constitutes the major component of plant biomass and could therefore be used as a raw material for food, clothing and energy. The model plant *Arabidopsis thaliana* can be used to study the SCW in cells of vascular bundles within the floral stem (Strabala and MacMillan, 2013). Anatomically, the vascular bundles are composed of phloem and xylem tissues and represent a central hub through which most biological compounds are transmitted to their site of use. Phloem tissue - composed of phloem parenchyma cells, companion cells and sieve elements (SE) - is involved in the transport of multiple compounds such as sugars, amino acids, proteins and mRNA (Le Hir *et al*., 2008; Zhang and Turgeon, 2018). A thickening of the phloem SE cell wall has been reported in Arabidopsis, and it was suggested that this cell wall is composed of pectic polysaccharides (Freshour *et al*., 1996). On the other hand, mature xylem tissue - responsible for structural support as well as the transportation of water and solutes - is composed of xylem tracheary elements (xylem vessels), xylary fibers and xylem parenchyma cells (Schuetz *et al*., 2012), and is characterized by the presence of thick SCWs. Recently, researchers used high-throughput immunolabelling of the major cell-wall glycan epitopes to cluster floral stem tissues according to their cell wall composition, with the results revealing a tissue-specific pattern for these glycan (Hall *et al*., 2013). However, we still lack precise information about cell wall composition at the cellular level. Among the tools that can provide spatial resolution at such a level, vibrational microspectroscopy approaches boast several strong advantages.

Vibrational spectroscopy techniques (e.g. Fourier-transformed infrared spectroscopy (FTIR), Raman spectroscopy) have been used extensively in plant research to decipher the cell wall composition in an organ-specific manner (Largo-Gosens *et al*., 2014). These complementary techniques offer many advantages among which the spatial resolution, the non-destruction of the samples and the cost-efficiency. They are also state-of-art techniques to study the plant lignocellulosic biomass (i.e. biosynthesis, degradation and valorization) that is of increasing interest due to its pivotal role for human and animal (i.e. food, energy, clothing, building material) (Gierlinger, 2017). Classically, FTIR microspectroscopy uses a thermal source to identify differences in the cell wall composition of wild-type and mutant of various plant species with a spatial resolution of approximately 30-50 μm (Sibout *et al*., 2005; Lefebvre *et al*., 2011; Le Hir *et al*., 2015). The coupling of a focal plan array (FPA) detector to a conventional FTIR microscope allows researchers to obtain structural information at the cellular level (Gorzsás *et al*., 2011; Ohman *et al*., 2013). Another powerful modification is the use of a synchrotron IR light source (SR-FTIR), which enables the collection of IR spectra at higher spatial resolution (i.e. cellular level) due to light that is at least 100 times brighter than that of a thermal source. However, this possibility has seen limited use in the plant biology field (Vijayan *et al*., 2015). In addition to IR microspectroscopy, Raman microspectroscopy is commonly used to study plant cell wall composition (Gierlinger *et al*., 2012; Gierlinger, 2017) including that of xylem cell types (i.e. *Arabidopsis thaliana* (Prats Mateu *et al*., 2016); *Cucumis sativus* (Zeise *et al*., 2018); *Malus sp.:* (Horbens *et al*., 2014); *Populus sp*. (De Meester *et al*., 2017; Jin *et al*., 2018); *Picea sp*. (Agarwal, 2006; Hänninen *et al*., 2011); *Pinus sp*. (Hänninen *et al*., 2011). To a lesser extent, the combined use of both techniques (e.g. FTIR and Raman microspectrocopy) has been proven efficient and complementary in studying the xylem cell wall composition in tree species such as Poplar (Özparpucu *et al*., 2017, 2018) and *Pinus radiata* (Zhang *et al*., 2017).

Regarding the plant model *Arabidopsis thaliana*, only a few studies have investigated the compositions of cell walls in the floral stem (Schmidt *et al*., 2010; Prats Mateu *et al*., 2016) despite the important physiological role of this plant organ. The floral stem, which supports the flowers and the fruits, is also a major contributor to lifetime carbon gain (Earley *et al*., 2009), representing 40% of a plant’s total biomass. The precise characterization of the cell wall compositions of different vascular cell types within the floral stem is necessary to better understand cell wall complexity within such tissues.

In addition to the identification of differences in plant cell wall composition at the tissue and cell levels, there are many unanswered questions regarding the modalities of sugar allocation, which directly influence the supply of carbohydrate skeletons required for SCW formation. We have previously focused on identifying the carbohydrate components underlying xylem secondary cell wall formation in *Arabidopsis thaliana*. By using conventional FTIR microspectroscopy, we identified cell wall modifications in the xylem of Arabidopsis double mutant defective in the expression of sugar facilitators encoding genes *SWEET11* and *SWEET12* (Le Hir *et al*., 2015). Both genes encode proteins that transport sugars (sucrose, glucose or fructose) along the concentration gradient (Chen *et al*., 2012; Le Hir *et al*., 2015) and, as such, their disruption modifies cellulose and xylan acetylation in xylem cell walls within the floral stem (Le Hir *et al*., 2015). In that study, the FTIR spectra were acquired over a 30x30 μm target zone, which encompasses different cell types. Moreover, the use of conventional FTIR on “dry-fixed” floral stem sections did not allow the acquisition of spectra describing phloem tissue due to complete shrinkage of the tissue when the floral stem section is dry. As *SWEET11* and *SWEET12* are expressed in both the phloem and xylem, spectral data at the cellular level are needed to better understand how modifications of sugar homeostasis influence the cell wall composition of various cell types (Chen *et al*., 2012; Le Hir *et al*., 2015).

In the presented work, we chose to use SR-FTIR and Raman microspectroscopy in combination with a Classification and Regression Tree (CART) ‐based method to analyze spectra collected for phloem and xylem tissues from *Arabidopsis thaliana* floral stem sections of wild-type and *sweet11-1sweet12-1* double mutant plants (Fig. 1). Overall, we show that SR-FTIR can be successfully used to analyze spectra acquired from phloem tissue, as well as that changes in *SWEET11* and *SWEET12* expression affect phloem cell wall composition. Additionally, the application of the CART method on Raman spectra shows that xylem vessels and fibers can be distinguished by a combination of cellulose and hemicellulose Raman shifts. Finally, our results suggest that facilitated sugar transport modifications in xylem parenchyma cells lead to cell-specific defects.

**Fig. 1.**
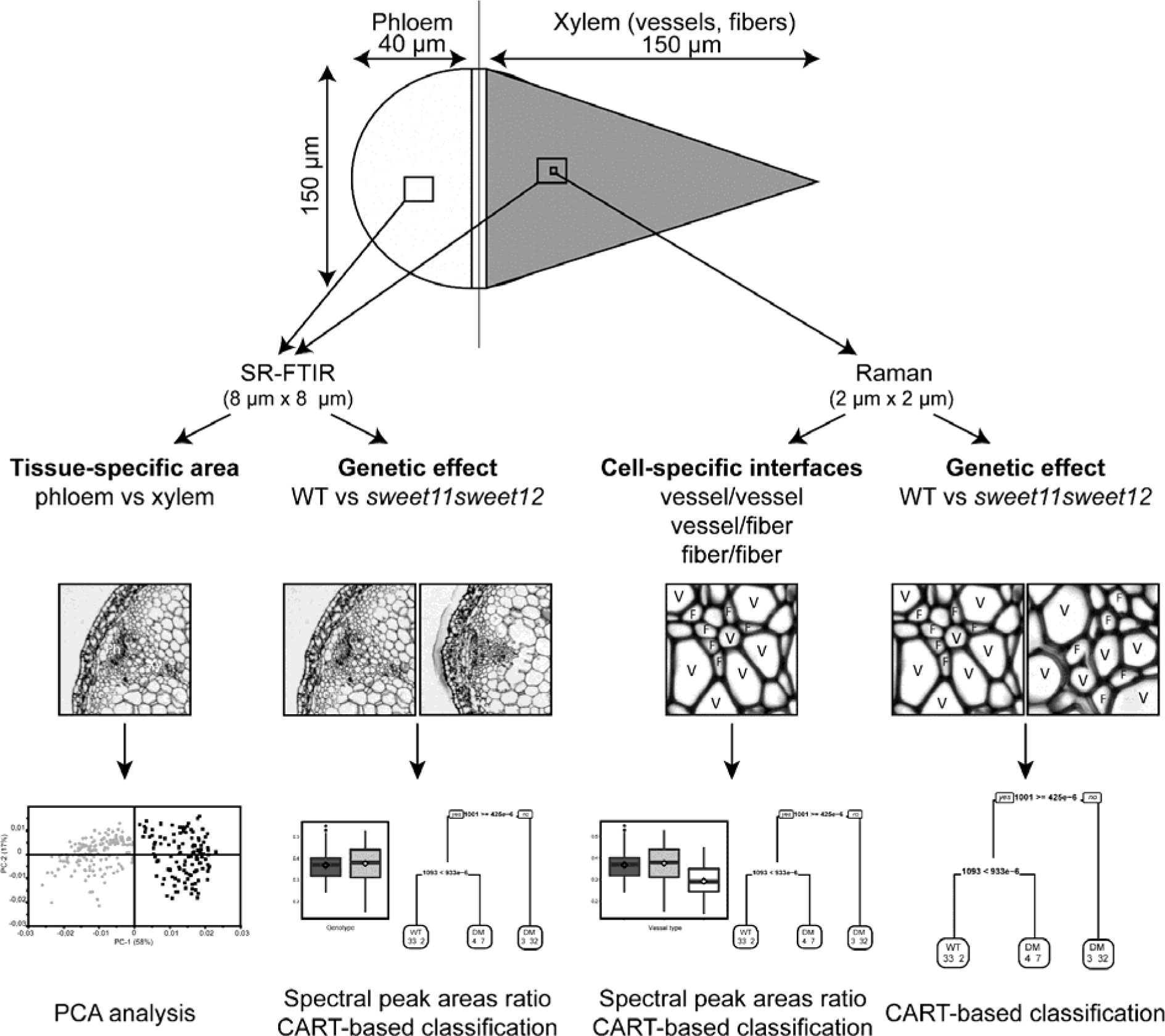
Workflow for the floral stem vascular system analysis by SR-FTIR and Raman spectroscopy. Schematic representation of a floral stem vascular bundle showing the phloem (light gray) and the xylem (dark gray) and the size of the acquisition zone for SR-FTIR and Raman analysis. SR-FTIR was used to obtain tissue-specific signature by PCA analysis of the spectra acquired on phloem and xylem cell walls of the WT plants. Additionally, a genetic effect was assessed by SR-FTIR by analyzing spectra of WT and mutant line with spectral peak areas ratios and CART-based classification. Then Raman microspectroscopy was used to obtain cell-specific interfaces signature on the different xylem cell types and to assess the genetic effect in a mutant line by spectral peak areas ratios and CART-based classification. F: fiber; Ph: phloem; SR-FTIR: synchrotron radiation fourier-transformed infrared spectroscopy; V: vessel; Xy: xylem.

## Materials and methods

### Plant material and growth conditions

Arabidopsis wild-type Col-0 line and *sweet11-1sweet12-1* (Le Hir *et al*., 2015) double mutants were grown in soil in a greenhouse for five weeks under long-day conditions (16 h photoperiod and 150 μE m^-2^ s^-1^ light intensity) at 22/15°C (day/night temperature) with 65% hygrometry. After five weeks of growth, the floral stem height is on average 25 cm for both genotypes. The first five centimeters of the basal part of floral stem were collected, fixed in 4% paraformaldehyde and embedded in paraffin. Sections with a thickness of 10 μm were deposited onto BaF2 windows and paraffin was removed using Histo-clear (National Diagnostics, Atlanta, GA). For SR-FTIR and RAMAN spectroscopy, four xylem/phloem poles from four plants representing each genotype were analyzed. Spectra were acquired on fully developed xylem vessels and fibers localized in the middle of the vascular bundle.

### Synchrotron radiation FTIR microspectroscopy (SR-FTIR)

Infrared spectra were recorded with a synchrotron source to provide better spatial resolution due to superior brightness (SOLEIL, SMIS beamline, Gif sur Yvette, France). The transmission spectra were collected on a NICOLET 5700 FT-IR spectrometer coupled to a Continuum XL microscope (Thermo Fisher Scientific, Waltham, MA) equipped with a 32X NA 0.65 objective as described in Guillon *et al*. (2011). All spectra were obtained in confocal mode to eliminate diffraction from surrounding cells using a double path single masking aperture size of 8 μm x 8 μm (Fig. S1B). The spectra were collected over the 1800-800 cm^-1^ infrared range at a spectral resolution of 4 cm^-1^ with 256 co-added scans for the background and sample spectra.

### Raman microspectroscopy

The Raman spectra were recorded using a DXR Raman Instrument (Thermo Fisher Scientific). Raman measurements were performed in a closed environment using a stabilized 532 nm laser as described in Zimmermann et al. (2015). A 100X NA 0.90 objective was used for focusing and collecting inelastically scattered Raman light, and allowed us to reach a spatial resolution of 2 μm x 2 μm (Fig. S1C). The acquisition points were set as the cell walls between xylem vessels (VV), between xylem vessels and fibers (VF) and between xylem fibers (FF). Overall the cell wall of 92 vessels and 126 fibers were analyzed in the WT floral stem while the cell wall of 115 vessels and 123 fibers were analyzed in the double mutant line. The system was operated in 25 μm aperture mode, which provided a spectral resolution of 2-4 cm^-1^. In order to decrease xylem cell wall autofluorescence, the samples were photobleached for two minutes before each acquisition. Sample spectra were acquired over an exposure time of 6 x 10 s using 512 scans (the same amount of scans were also performed for the background).

### Preprocessing of SR-FTIR and Raman spectra

Infrared spectra with extreme absorbance values, e.g. values less than 0.1 or above 1, were removed from the datasets so that saturation effects and errors due to holes in the tissue sections could be avoided. Spectra comparison between tissue or genotype was performed on baseline-corrected and area-normalized spectra. For Raman microspectroscopy, the spectra of different xylem cell types were smoothed by the Savitsky-Golay algorithm (3^rd^ order polynomial and nine-point filter). The spectra were then baseline-corrected by subtracting a linear baseline between 350-3500 cm^-1^ and area-normalized. Both SR-FTIR and Raman spectra were preprocessed using Unscrambler software (The Unscrambler, CAMO Process AS, Oslo, Norway).

### Univariate analysis of the SR-FTIR and Raman spectra

Peak area measurements were performed on baseline-corrected and area-normalized FTIR spectra in OMNIC 9.2.41 and TQ Analyst EZ 9.2.34 software (Thermo Scientific). In both cases, the baseline between the peak start and end marker was computer generated. The following peaks were measured for the SR-FTIR spectra: 930-1180 cm^-1^ for cellulose; 1695-1770 cm^-1^ for hemicellulose; and 1475-1520 cm^-1^ for lignin. Average spectra and boxplot representations were generated in R software (R Development Core Team, 2016) using the HyperSpec and Ggplot2 packages, respectively (Wickham, 2009; Beleites, 2012). Peak areas and their ratios were first checked for normality (Shapiro-Wilk test) and homoscedasticity (Levene test). Since neither of these criteria were fulfilled, an approximate (Monte Carlo) Fisher-Pitman permutation test was performed (non-parametric one-way ANOVA). Then, a pairwise comparison test, including the calculation of an adjusted *P*-value by the False Discovery Rate (FDR) method, was applied to assess the significance of observed differences. Statistical analysis was performed using the “coin”, and “RVAideMemoire” packages (Hothorn *et al*., 2008; Maxime, 2017) in R software (R Development Core Team, 2016).

### Multivariate statistical analysis of SR-FTIR and Raman spectra

Preprocessed and mean-centered spectra were first subjected to Principal Component Analysis (PCA). The PCA was carried out with three to seven principal components (PC) using the NIPALS algorithm, and full cross-validation was applied. Outliers identified using the Hotelling T2 method (95% multivariate confidence interval) and the residual versus leverage plot were removed from the dataset. Since SR-FTIR and Raman techniques produce large dataset and variables are highly correlated, a variable selection algorithm (CovSel) was applied prior to the Classification and regression tree (CART) technique. The CovSel algorithm enables variable selection based on global covariance across all the responses (Roger *et al*., 2011). Additionally, the CART technique can be used to select the variables that are most important to discriminating two factors (Berk, 2016). The CART-based model was first set up on a calibration dataset (representing 80% of the total spectra) and then validated on a validation dataset (representing 20% of the total spectra). A confusion table was then produced to validate the model. Model performance was evaluated using the following parameters: accuracy 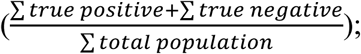 specificity 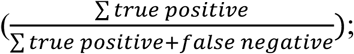 sensitivity 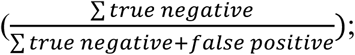 predictive positive value 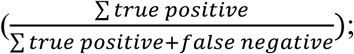 and negative predictive value 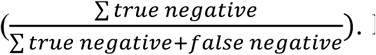. In the comparison of both genotypes, the true positive and true negative represent the number of wild-type or *sweet11-1sweet12-1* spectra that were correctly classified by the model. The false positive and false negative represent the number of wild-type or *sweet11-1sweet12-1* spectra that were incorrectly classified by the model. The multivariate analyses were performed in the ChemFlow interface within Galaxy (https://vm-chemflow.toulouse.inra.fr/).

## Results and discussion

### Synchrotron radiation FTIR (SR-FTIR) allows the identification of unique spectral fingerprints for the floral stem vascular tissues

In the Arabidopsis floral stem, vascular tissues are organized as a series of vascular bundles that are connected together by the interfascicular cambium (Fig. S1A). In each bundle, specialized conducting cells are subjected to high pressures to ensure sap flow, with hydrostatic pressure reaching upwards of 30 atmospheres in the sieve elements (Sjölund, 1997) while the xylem vessels are characterized by negative pressure. Cell-specific cell wall composition is a crucial part of conferring resistance to such pressure. A majority of the previous research has focused on deciphering xylem cell wall composition, while phloem tissue has received limited attention.

By harnessing the spatial resolution provided by Synchrotron light (8x8 μm acquisition zone), we acquired spectra for phloem and xylem tissues of the Arabidopsis wild-type floral stem (Fig. 1 and Fig. S1A). Spectra were baseline corrected, area-normalized and the average spectrum for each tissue was calculated and plotted (Fig. 2A). Principal component analysis (PCA) was used to identify potential spectral fingerprints of the floral stem vascular tissues, with the first two components explaining 54% and 8% of the total variance (Fig. 2B). Component 1 undoubtedly discriminates the phloem and xylem IR spectra (Fig. 2B). The loading plot of PC1 reveals that xylem cell walls are characterized by a set of bands corresponding to guaiacyl ring breathing with carbonyl stretching (1269 cm^-1^) (Kubo and Kadla, 2005), -C-H-deformation in the guaiacyl ring with -C-O-deformation in the primary alcohol (1030 cm^-1^) (Kubo and Kadla, 2005) and ‐C-C-linkage of G-condensed units (1060 cm^-1^). This suggests that xylem cell walls are mainly composed of G-type lignin (Fig. 2C and Table 1), and these results are in agreement with previous observations from *Arabidopsis thaliana* that showed that G-type lignin is responsible for the extra-thickening of the xylem vessel cell wall (Schuetz *et al*., 2012). Additionally, the loading plot highlights several wavenumbers (1045, 1369, 1230-1235, 1743, 1245, 1735-1740 cm^-1^) that are related to hemicellulose enrichment in xylem cell walls, as has been previously reported for the Arabidopsis floral stem (Table 1) (Sibout *et al*., 2005; Ohman *et al*., 2013). However, some of these bands may partly overlap with the lignin bands (1236, 1371 and 1736 cm^-1^) (Faix, 1991; Özparpucu *et al*., 2018).

**Fig. 2.**
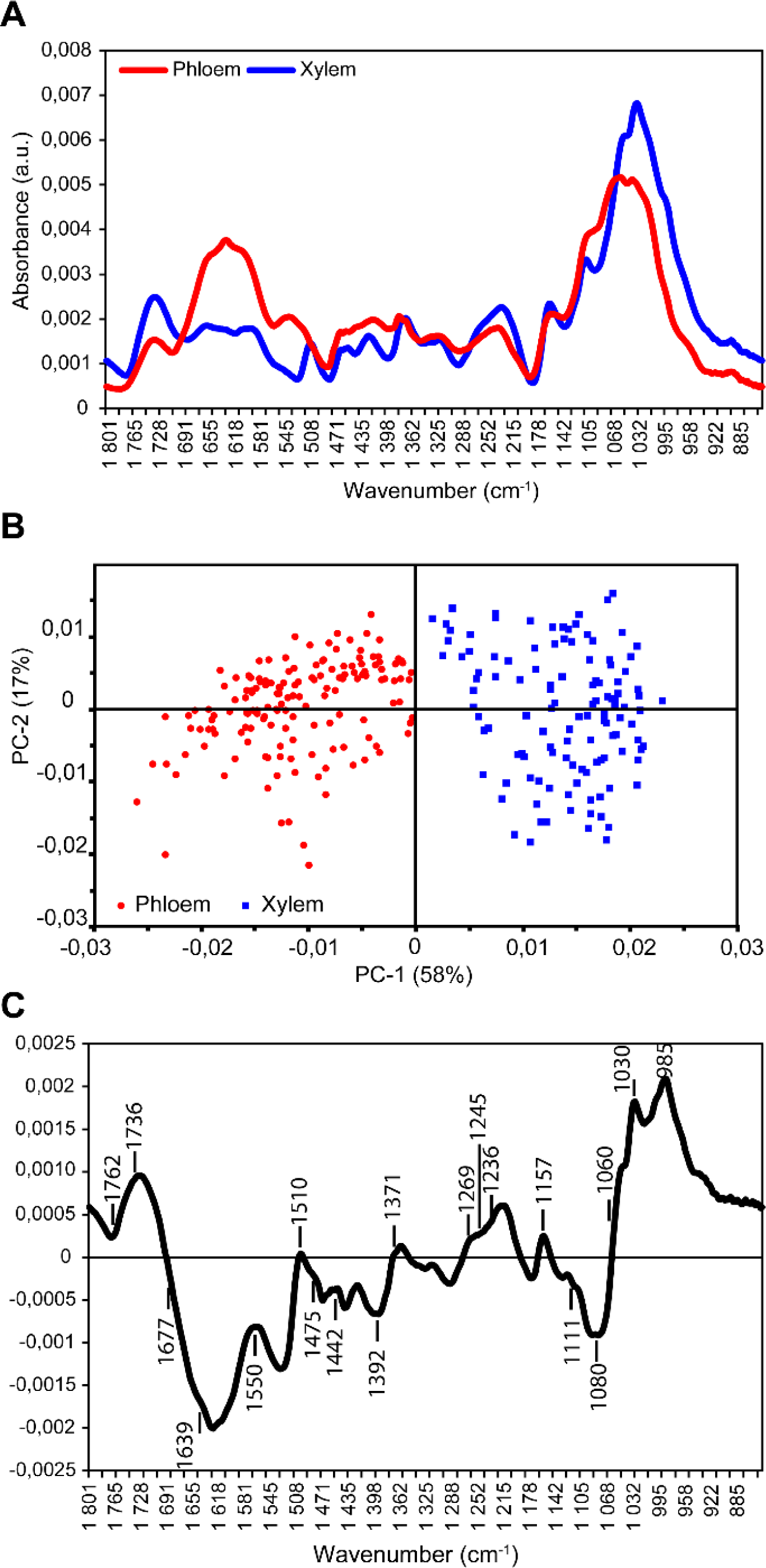
Principal Component Analysis (PCA) of infrared spectra obtained from phloem and xylem cells of Arabidopsis wild-type floral stem. (A) Average spectra for wild-type phloem (red line) and xylem (blue line) tissues obtained by SR-FTIR microscopy. Spectra were baseline-corrected and area-normalized in the range of 1800-850 cm^-1^. (B) Comparison of phloem and xylem cell wall composition by multivariate analysis. Spectra from xylem (blue boxes) or phloem (red circles) cell walls were compared. A scoreplot based on PC1 and PC2 from the Principal Component Analysis (PCA) shows that phloem and xylem cell wall spectral signatures can be differentiated. (C) The corresponding loading plot of the PC1 axis is presented.

**Table 1.** Assignment of the infrared wavenumbers found to differentiate xylem and phloem tissues of the wild-type Arabidopsis floral stem.

| Tissue | FTIR wavenumber (cm<sup>-1</sup>) | Assignment | Polymer |
| --- | --- | --- | --- |
| Xylem | 1510 | G-type lignin (Faix, 1991) | Lignin |
|  | 1595 | G-type lignin (Faix, 1991) | Lignin |
|  | 1269 | Guaiacyl ring breathing with carbonyl stretching (Kubo and Kadla, 2005) | Lignin |
|  | 1030 | C-H deformation in guaiacyl with C-O deformation in the primary alcohol (Kubo and Kadla, 2005) | Lignin |
|  | 1060 | C-C linkage of G condensed unit (Sibout *et al.*, 2005) | Lignin |
|  | 1045 | C-O-C contribution of xylan (Brown *et al.*, 2009) | Hemicellulose |
|  | 1369 | Deformation of the C-H linkages in the methyl group of *O*-acetyl moieties (Mohebbi, 2010) | Hemicellulose |
|  | 1230-1235 | C=O/C-O linkages stretching vibrations (Mohebbi, 2010) | Hemicellulose |
|  | 1743 | Stretching of the free carbonyl group (Owen and Thomas, 1989) | Hemicellulose |
|  | 1245 | C-O stretch (Faix, 1991) | Hemicellulose |
|  | 1735-1740 | C=O stretching in glucuronic acid (xylan) (Marchessault, 1962; Marchessault and Liang, 1962) | Hemicellulose |
|  | 1762 | Esterified pectins (Kačuráková *et al.*, 2002) | Pectins |
| Phloem | 1639 | COOH group (Mouille *et al.*, 2006) | Acidic pectins |
|  | 1677 | COOH group (Mouille *et al.*, 2006) | Acidic pectins |
|  | 1157 | C-O-C linkages of cellulose (Kačuráková *et al.*, 2002) | Cellulose |
|  | 1442 | To be assigned (Mouille *et al.*, 2003) | Related to primary cell wall |
|  | 1111 | In plane ring stretching (Bekiaris *et al.*, 2015) | Cellulose |
|  | 1712 | C=O stretch (Kačuráková *et al.*, 2002) | Pectins |
|  | 978 | Xylan-type polysaccharides (Brown *et al.*, 2005) | Hemicellulose |
|  | 958 | Sugar ring vibrations (Kačuráková *et al.*, 2002) | Pectins |
|  | 1475 | To be assigned | To be assigned |
|  | 1550 | Carboxylates (Mouille *et al.*, 2006) | Cellulose |
|  | 1774 | To be assigned | Esterified pectins |

Interestingly, wavenumbers associated with pectic polysaccharides (1245 and 1762 cm^-1^) were also found to be more descriptive of xylem cell walls than of phloem cell walls (Fig. 2C and Table 1), even if pectins are not an abundant component of secondary cell walls. However, pectin methylesterification appears to be a prerequisite for the lignin modification that occurs during secondary cell wall deposition in xylem cells (Pelloux *et al*., 2007). In addition, several Arabidopsis mutants that are deficient in various pectins have been shown to present defects in secondary cell wall formation (Persson *et al*., 2007; Lefebvre *et al*., 2011).

Regarding phloem cell wall composition, the loading plot of PC1 reveals numerous wavenumbers related to pectic polysaccharides, cellulose and hemicelluloses (Fig. 2C and Table 1). For instance, the bands at 1639 cm^-1^ and at 1677 cm^-1^ are characteristic of the -COOH-group of acidic pectins present in the primary cell wall (Mouille *et al*., 2006), while wavenumbers at 1111, 1157 and 1550 cm^-1^ describe cellulose polymers (Table 1). Additionally, wavenumbers at 1442 and 1475 cm^-1^ were unique for phloem cell walls. The 1442 cm^-1^ band has been previously reported to describe the hypocotyl primary cell wall of a cellulose-deficient mutant, but the functional group that it represents still needs to be verified (Mouille *et al*., 2003). Overall, these results suggest that the cell wall composition of phloem tissue, including phloem parenchyma cells, companion cells and sieve elements, is more closely related to primary cell wall composition even if cell wall thickening is commonly observed in sieve elements (SEs) (Esau and Cheadle, 1958). The nature of this thickening has not yet been completely clarified, but the current evidence favors a pectin-based composition (Freshour *et al*., 1996; Torode *et al*., 2018). The marker IR bands we identified in our tissue samples support these findings. It was initially surprising that pectins, which have been traditionally related to cellular expansion, can be found in a tissue that experiences high pressure during sap flow. The recent characterization of an antibody against branched pectic galactan that specifically binds to the cell walls of SEs led the authors to suggest that the role of pectin in these cell walls could be the maintenance of elastic properties required for withstanding high turgor pressure (Torode *et al*., 2018). Additionally, the application of atomic force microscopy (AFM) has shown that the mechanical properties of phloem SE cell walls differ from those of cells from the surrounding tissue with an higher elasticity (Johnson, 2018; Torode *et al*., 2018). The recent development of the nano-IR techniques that combine AFM and Synchrotron IR light (Pereira *et al*., 2018) will open further possibilities for exploring the cell wall heterogeneity that exists among different vascular cell types. Further the implementation of microfluidic infrared spectroscopy to study plant sample in a liquid environment represents an important progress (Devaux *et al*., 2018) especially to study plant tissue that are not accessible anymore after the dehydration process, usually required for IR analysis, such as the phloem tissue.

### Phloem cell wall composition is impaired in the double mutant sweet11-1sweet12-1

In addition to presence in the xylem tissue, *SWEET11* and *SWEET12* expression has also been detected in phloem tissue, with the signal arising from phloem parenchyma cells (Chen *et al*., 2012; Le Hir *et al*., 2015). Since the double mutant *sweet11-1sweet12-1* shows defects in xylem cell wall formation, we analyzed whether similar defects in cell wall formation could be observed in phloem cells. Therefore, phloem and xylem IR spectra were recorded from the double mutant *sweet11-1sweet12-1* floral stem and compared to spectra acquired from the wild-type floral stem (Fig. 1). The average spectra for wild-type (WT) and double mutant (DM) tissues were compared after baseline correction and area normalization over the 1800-850 cm^-1^ range (Supplementary Fig S2A and S2E). Next, the areas under the cellulose (C-O and C-C stretching) (930-1180 cm^-1^), the hemicellulose (1695-1770 cm^-1^) and/or lignin (1475-1520 cm^-1^) peaks, along with their respective ratios, were measured (Fig. 3 and Supplementary Fig. S2B-D and S2F-G). A significant decrease in the lignin peak area of the xylem tissue was measured between both genotypes (Supplementary Fig. S2D), while a significant increase in the cellulose and the hemicelluloses peak areas was observed in xylem cell walls in the double mutant line compared to the wild-type (Supplementary Fig. S2B and C). The phloem tissue analysis showed that there was no significant difference in cellulose between the two genotypes (Supplementary Fig. S2F), while the hemicelluloses peak area in the double mutant line was significantly greater that what was observed in the wild-type (Supplementary Fig. S2G). More precisely, while the xylem cell walls of both genotypes showed similar cellulose/hemicellulose ratios (Fig. 3A), the phloem cell walls of the double mutant demonstrated a disequilibrium in the cellulose/hemicellulose ratio (Fig. 3B). Therefore, we show that in addition to affecting xylem cell wall composition, mutations in both *SWEET11* and *SWEET12* genes also impact phloem cell wall composition. However, these mutations seem to only affect the hemicellulose composition of phloem cell walls.

**Fig. 3.**
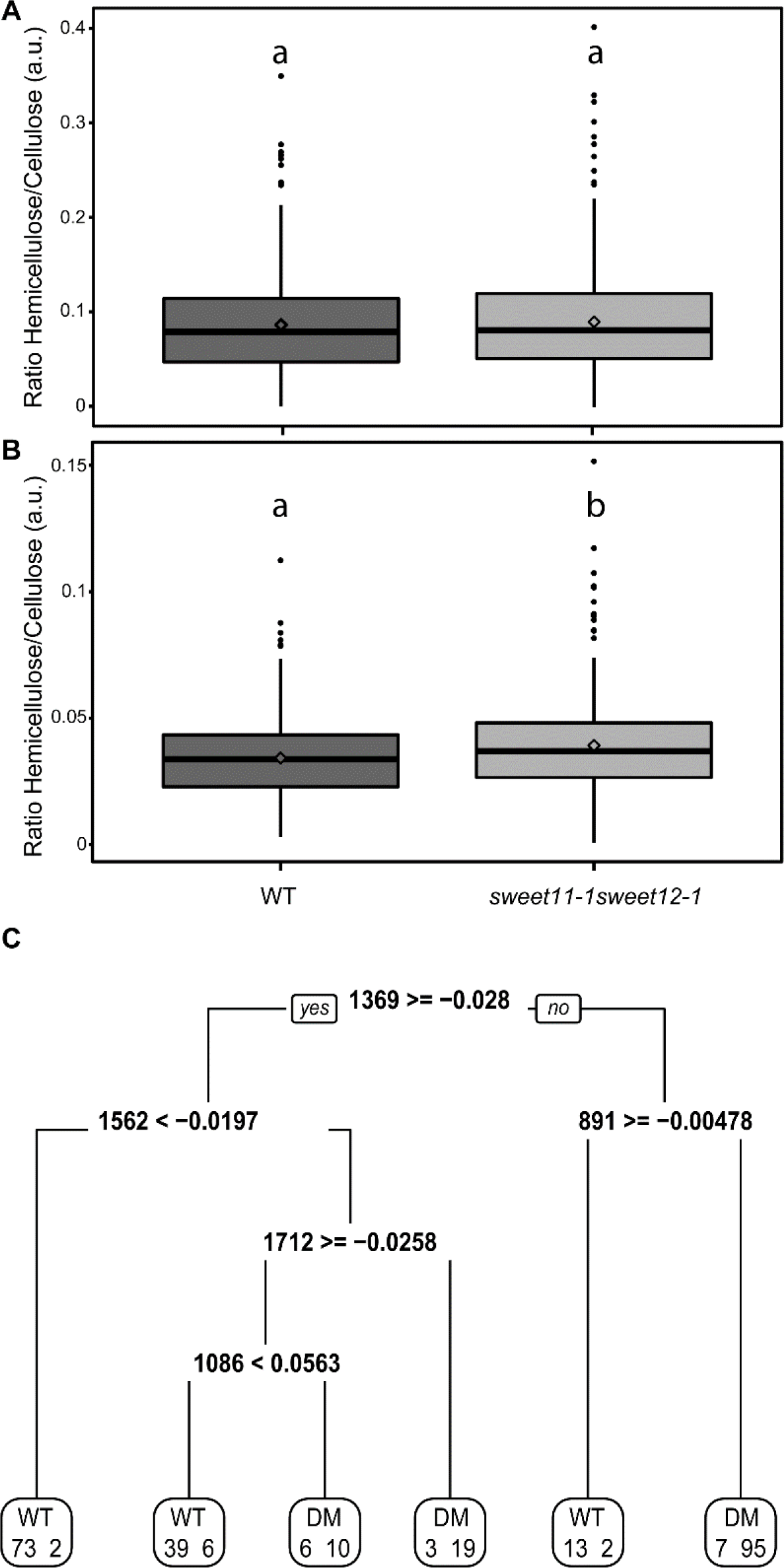
Comparative analysis of wild-type and *sweet11-1sweet12-1* xylem or phloem SR-FTIR spectra. (A-B) Boxplot representation of the hemicellulose/cellulose ratio of xylem (A) and phloem (B) spectra. For the xylem spectra, the box and whisker plots represent values from 521 and 494 individual spectra of wild-type and *sweet11-1sweet12-1* lines, respectively. For the phloem spectra, the box and whisker plots represent the values from 314 and 311 individual spectra of wild-type and *sweet11-1sweet12-1* lines, respectively. The diamonds represent mean values, lines represent median values, the tops and bottoms of the boxes represent the first and third quartiles, respectively, and whisker extremities represent maximum and minimum data points. The black dots are the outliers. Letters above the boxes indicate groups with significant differences as determined by an approximate Fisher-Pitman permutation test and a pairwise comparison test (*P* < 0.05). a.u.: arbitrary unit. (C) The classification tree has been generated by the CART method after ten-fold cross-validation of the calibration dataset model, which was built using the 850-1800 cm^-1^ range of phloem spectra. The binary classification tree is composed of five classifiers and 6 terminal subgroups. The decision-making process involves the evaluation of if-then rules of each node from top to bottom, which eventually reaches a terminal node with the designated class outcome (WT: wild-type and DM: *sweet11-1sweet12-1*). The numbers in each terminal subgroup represent numbers of either WT or DM spectra.

In order to further identify wavenumbers that could be specifically associated with the *sweet11-1sweet12-1* double mutant, we applied a CART analysis procedure. For this purpose, our original dataset was split into calibration (80% of the total dataset) and validation (20% of the total dataset) datasets, after which the CovSel algorithm was applied to the calibration dataset to identify the 10 wavenumbers with maximum covariance (Roger *et al*., 2011). The CART tree resulting from the analysis shows that, out of the 10 selected wavenumbers, only five IR wavenumbers - at 891, 1086, 1369, 1562 and 1712 cm^-1^ – can be used to distinguish between the wild-type and *sweet11-1sweet12-1* phloem spectra (Fig. 3C). To evaluate the performance of this analysis, the CART model obtained from the calibration dataset was used as an input and applied on the validation dataset. Table 2 summarizes the results of both genotypes for the model calibration (after ten-fold cross-validation) and validation datasets. When applied on the calibration dataset, the CART model correctly classified 88.6% of the wild-type spectra (specificity) and 92.5% (sensitivity) of the double mutant spectra. When applied on the validation dataset, the model correctly classified 82.7% (specificity) and 79.5% (sensitivity) of the WT and DM spectra, respectively. Moreover, the predictive positive value (PPV) of the validation model was calculated to be 75%, which means that most of the identified WT spectra are not false positives (Table 2). On the other hand, the model’s negative predictive value (NPV) was determined to be 86.1%, which means that a majority of the identified *sweet11-sweet12-1* DM spectra are not false positives (Table 2). Overall, the CART model was able to accurately predict 80.8% of the spectra present in the validation dataset (Table 2). In this way, the CART model produced using the calibration dataset can discriminate both genotypes based only on the analysis of five major FTIR wavenumbers. Among these marker wavenumbers, the 891 cm^-1^ wavenumber can be linked to the cellulose fingerprint region (Kačuráková *et al*., 2002) while the 1086 cm^-1^ and 1369 cm^-1^ wavenumbers can be assigned to hemicelluloses (Robin *et al*., 2003; Brown *et al*., 2005). Additionally, the 1712 cm^-1^ wavenumber could be related to carboxylic acid residues found in polygalacturonic acid (Pawar *et al*., 2013). The remaining wavenumber, 1562 cm^-1^, still needs to be assigned to a cell wall compound. Interestingly, we previously found that the 1369 cm^-1^ wavenumber (this work and (Le Hir *et al*., 2015)) can differentiate WT xylem cell walls from the cell walls of the *sweet11-sweet12-1* double mutant. This wavenumber is related to the deformation of C-H linkages in the methyl group of O-acetyl moieties and could thus represent differences in xylan acetylation (Mohebby, 2010). Therefore, the presented results suggest that sugar homeostasis modifications in plant vascular tissue predominantly influence the cellulose and/or xylan composition of cell walls regardless of cell type.

**Table 2.**
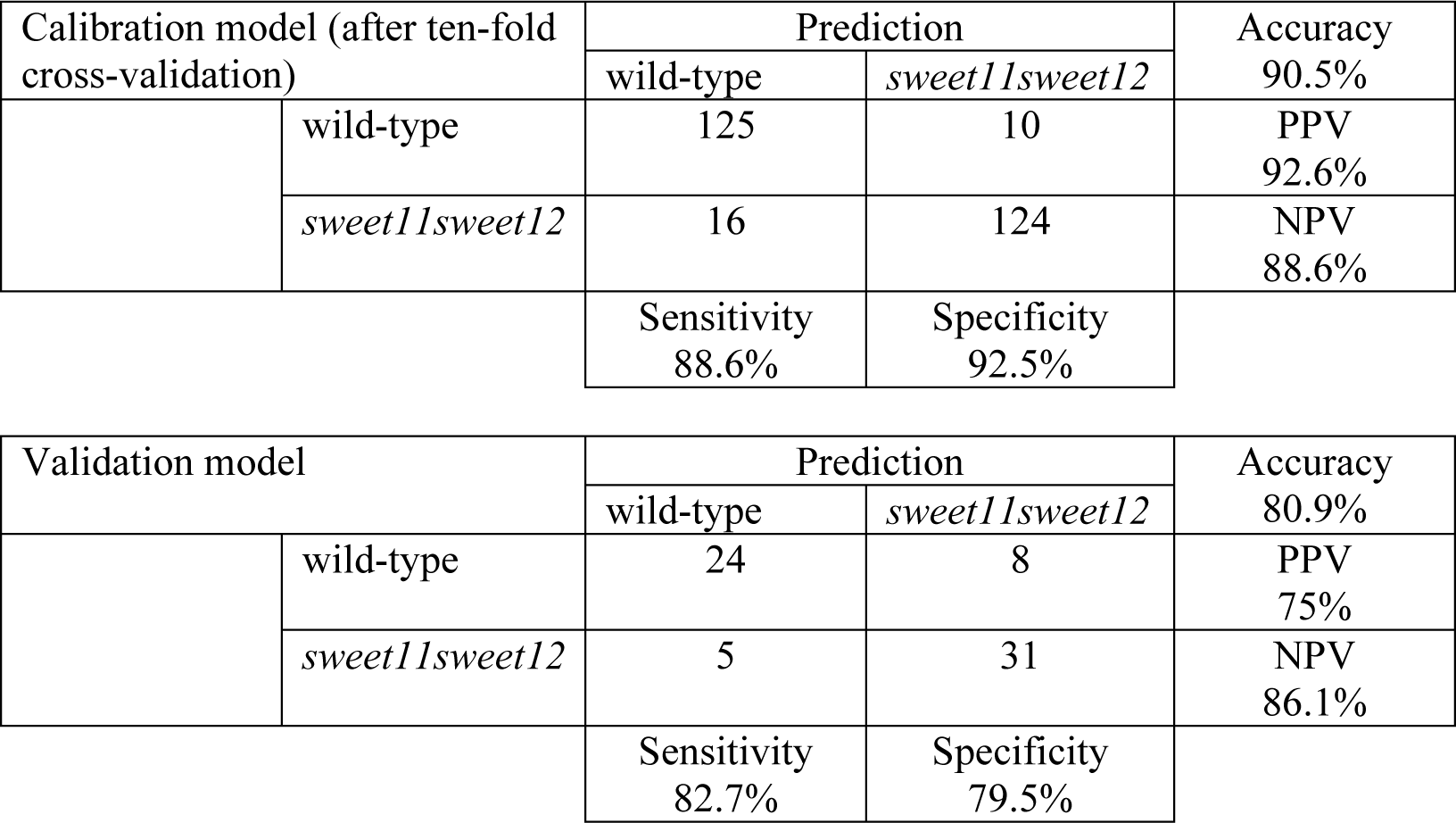
Classification results for using FTIR wavenumbers to predict which genotype a phloem tissue sample represents, with the model calibration dataset (80% of total dataset) using a ten-fold cross-validation method, and the validation dataset (20% of total dataset) using a CART-based algorithm. NPV: negative predictive value, PPV: positive predictive value.

We previously postulated that the maintenance of sugar homeostasis among the xylem parenchyma cells and xylem vessels/fibers influenced the production of a normal cell wall (Le Hir *et al*., 2015). Here, we show that it also constitutes a limiting step for the formation of phloem cell walls. Since *SWEET11* and *SWEET12* are expressed in the phloem and xylem parenchyma cells and participate in sugar influx or efflux across the plasma membrane (Chen *et al*., 2012; Le Hir *et al*., 2015), our data suggest that *SWEET11* and *SWEET12* are crucial for cell wall formation in vascular parenchyma cells. Interestingly, recent research has also found vascular parenchyma cells to be crucial in the supply of monolignols to developing xylem vessels (Smith *et al*., 2017). Therefore, one could postulate that sugar (sucrose and/or hexoses) movement across a gradient, mediated by SWEET11 and/or SWEET12, could also occur between vascular parenchyma cells and other developing vascular cells to drive cell wall formation.

### Identification of new Raman shift markers that describe the composition of cell walls between different xylem cell types in the wild-type Arabidopsis floral stem

Xylem secondary cell wall formation constitutes a large pool of the plant’s total biomass. For example, the xylem vessels and fibers are surrounded by a thick SCW that is 80% cellulose and hemicelluloses and 20% lignin (Marriott *et al*., 2016). When the Arabidopsis floral stem is considered at the cellular level, xylem vessels and fibers demonstrate heterogeneous cell wall composition due to differences in the lignin monomer(s) with which the cell wall is enriched (Schuetz *et al*., 2012). Unfortunately, we still lack a complete description of the polysaccharide composition of xylem vessel and fiber cell walls. We leveraged the spatial resolution provided by Raman microspectroscopy to precisely characterize the composition of cell walls between xylem vessels (VV), between xylem vessels and xylem fibers (VF) and between xylem fibers (FF) in the Arabidopsis wild-type floral stem (Fig. 1, Fig. S1C, Fig. 4 and Supplementary Fig. 3).

**Fig. 4.**
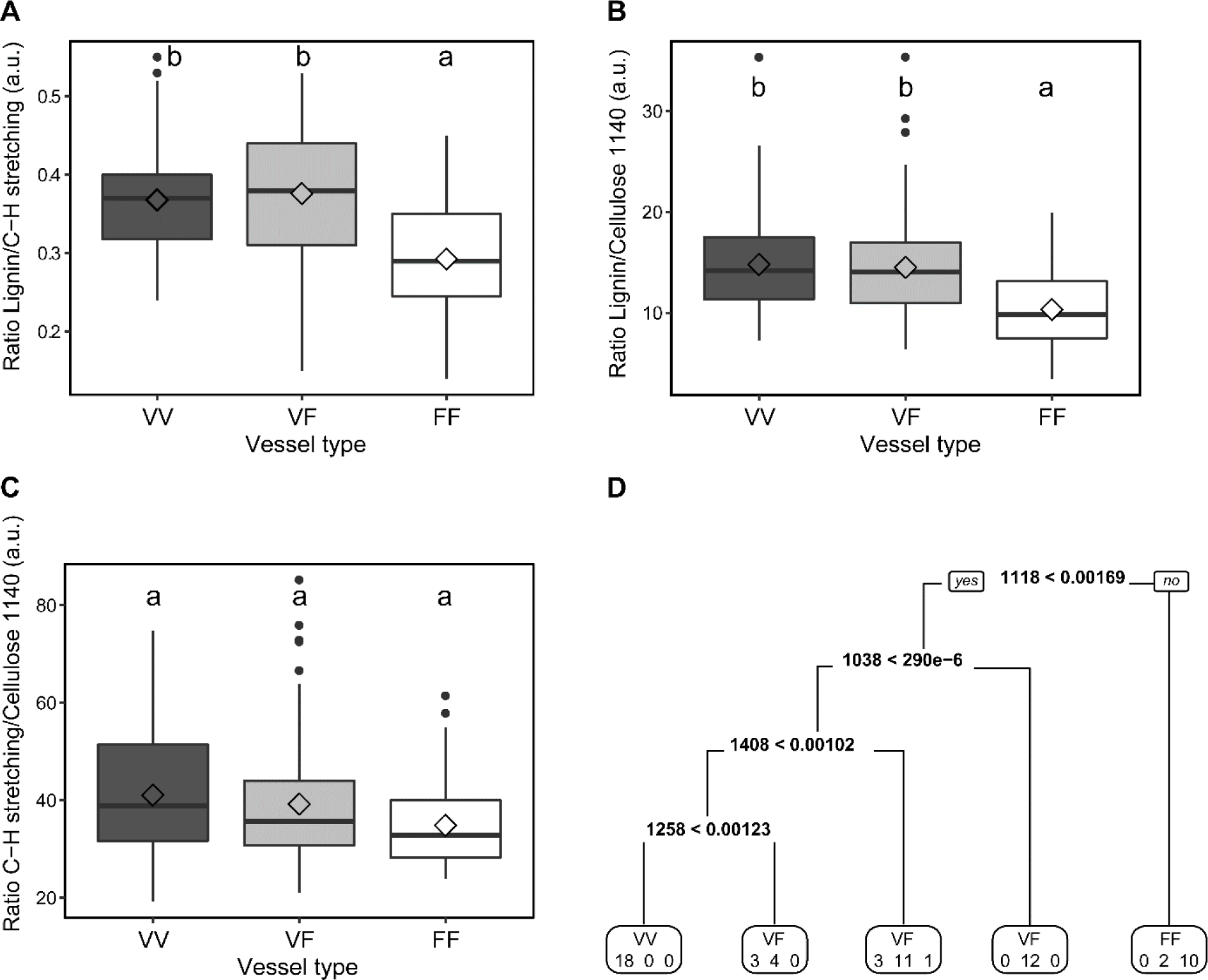
Raman spectra analysis of the different xylem cell types in wild-type plants. (A-C) Boxplot representation of the lignin/C-H stretching band ratio (A), lignin/C-O and C-C bond stretching ratio (B) and C-H stretching band/C-O and C-C bond stretching ratio (C) in secondary cell walls between different xylem cell types. The box and whisker plots represent values from 51, 139 and 119 individual spectra of VV, VF and FF, respectively. The diamonds represent mean values, lines represent median values, the tops and bottoms of the boxes represent the first and third quartiles, respectively, and whisker extremities represent maximum and minimum data points. The black dots are the outliers. Letters above the boxes indicate groups with significant differences as determined by an approximate Fisher-Pitman permutation test and a pairwise comparison test (*P* < 0.05). a.u.: arbitrary unit. (F) The classification tree has been generated by the CART method after ten-fold cross-validation of the calibration dataset model, which was built using the 1000-1800 cm^-1^ range from the Raman spectra of different xylem cell types. The binary classification tree is composed of four classifiers and 5 terminal subgroups. The decision-making process involves the evaluation of if-then rules of each node from top to bottom, which eventually reaches a terminal node with the designated class outcome (VV: vessel/vessel cell wall, VF: vessel/fiber cell wall and FF: fiber/fiber cell wall). The numbers in each terminal subgroup stand for the number of VV, VF or FF spectra.

The average Raman spectra for the various xylem cell types show that xylem fiber cell wall composition differs in comparison to what was observed in the other two cell types (Supplementary Fig. S3A). To test whether these differences were statistically significant, we calculated ratios of spectral peaks areas from already known Raman shift markers, namely, from 2775 to 3125 cm^-1^ (the composite C-H stretching bands comprising cellulose and hemicelluloses), from 1550 to 1700 cm^-1^ (lignin Raman shift) and from 1080 to 1140 cm^-1^ (C-O and C-C bond stretches of cellulose) (Schmidt *et al*., 2010; Agarwal, 2014) (Fig. 4). Based on these measurements, the VV and VF cell walls in wild-type Arabidopsis plants could not be statistically distinguished (Fig. 4A-C). However, the ratio of lignin to C-H bonds as well as the ratio of lignin to C-O bonds can significantly discriminate the cell walls between xylem fibers (FF) from those between xylem vessels (VV) and xylem vessels and fibers (VF) (Fig. 4A and 4B). There were no significant differences in the ratio of C-H bonds to C-O bonds between cell types (Fig. 4C). Therefore, the observed differences between cell walls between VV, VF and FF can mainly be attributed to a lower intensity of the aromatic ring stretching vibration (1598 cm^-1^) in the cell walls between xylem fibers (Fig. S3A) (Özparpucu *et al*., 2017).

To further identify Raman shifts associated with different xylem cell types, a CART-based classification method was applied on the 1000-1800 cm^-1^ Raman shift range, which includes the predominant constituents of the xylem cell wall (Prats Mateu *et al*., 2016; Özparpucu *et al*., 2017). The CART model was built on the calibration dataset (80% of the total dataset), without a variable selection step, and the resulting classification tree shows that only four Raman shifts are sufficient to distinguish the three different cell wall types (Fig. 4D). The overall accuracy of the model produced from the calibration dataset was 85.9%, with good prediction values (PPV) of 100%, 77.7% and 83.3% for the VV, VF and FF groups, respectively (Table 3). This model was then applied to the validation dataset (20% of the total dataset), and showed an accuracy value of 80%, which is close to that of the calibration dataset (Table 3). Even though the predictive sensitivity for spectra between adjacent xylem vessels (VV) was low, with only 50% of spectra correctly classified as VV spectra (Table 3), the PPVs for VF and FF spectra (87.5 and 100%, respectively) were good (Table 3). Therefore, our CART model can be used to distinguish VF and FF spectra, but the results should be interpreted with caution in the case of VV spectra. Nevertheless, our data show that the 1038, 1118, 1408 and 1258 cm^-1^ Raman shifts can be used to discriminate most of the different xylem cell wall types in wild-type Arabidopsis plants. Interestingly, the band around 1038 cm^-1^ was reported to describe C-O stretching of mannan oligosaccharides (Maru *et al*., 2015) while the Raman shift around 1256 cm^-1^ has been linked to hemicelluloses (Gierlinger *et al*., 2008). The bands at 1121 cm^-1^ (symmetric υ(COC) glycosidic bond) and 1408 cm^-1^ (δ(CH_2_) region) have also been assigned to cellulose (Edwards *et al*., 1997; Chylinska *et al*., 2014).

**Table 3.**
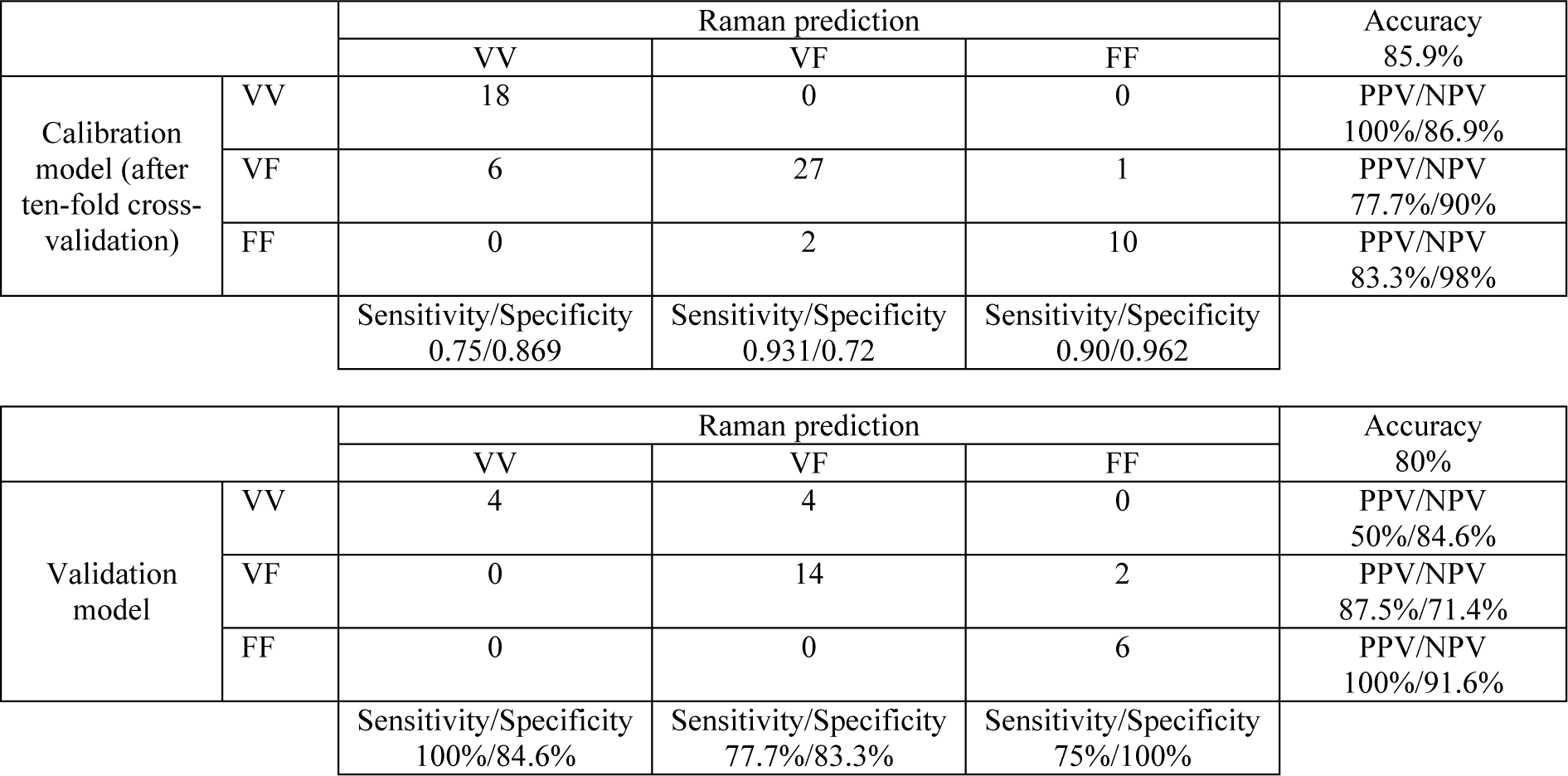
Classification results for using Raman shifts to predict different xylem cell types, with the model calibration dataset (80% of the total dataset) using a ten-fold cross-validation method, and the validation dataset (20% of the total dataset) using a CART-based algorithm. For the calculation of the different parameters, one cell type was compared to the two others. FF: cell wall between two xylem fibers, NPV: negative predictive value, PPV: positive predictive value, VF: cell wall between xylem vessel and fiber, VV: cell wall between two xylem vessels.

|  |  | Raman prediction |  |  | Accuracy |
| --- | --- | --- | --- | --- | --- |
|  |  | VV | VF | FF | 85.9% |
| Calibration<br>model (after<br>ten-fold cross-validation) | VV | 18 | 0 | 0 | PPV/NPV<br>100%/86.9% |
|  | VF | 6 | 27 | 1 | PPV/NPV<br>77.7%/90% |
|  | FF | 0 | 2 | 10 | PPV/NPV<br>83.3%/98% |
|  |  | Sensitivity/Specificity<br>0.75/0.869 | Sensitivity/Specificity<br>0.931/0.72 | Sensitivity/Specificity<br>0.90/0.962 |  |
|  |  | VV | VF | FF | 80% |
| Validation<br>model | VV | 4 | 4 | 0 | PPV/NPV<br>50%/84.6% |
|  | VF | 0 | 14 | 2 | PPV/NPV<br>87.5%/71.4% |
|  | FF | 0 | 0 | 6 | PPV/NPV<br>100%/91.6% |
|  |  | Sensitivity/Specificity<br>100%/84.6% | Sensitivity/Specificity<br>77.7%/83.3% | Sensitivity/Specificity<br>75%/100% |  |

Earlier studies in Arabidopsis have clearly established that the cell walls of xylem interfascicular fibers and xylem vessels differ in terms of their lignin monomer composition (Schuetz *et al*., 2012). Additionally, results from Poplar studies suggest that the cell wall composition of xylem fibers is an intermediate between that of xylem interfasicular fibers and xylem vessels (Gorzsás *et al*., 2011). ToF-SIMS has previously been applied to measure differences in the S/G ratio between xylem fibers and vessels in *Populus* (Tolbert *et al*., 2016). Our work shows that a combination of Raman shifts assigned to cellulose and hemicelluloses can also distinguish xylem cell types. This is in agreement with previous research, as the immunolabelling of mannan epitopes (LM10 and LM11 antibodies) in the Arabidopsis floral stem revealed a higher signal intensity in xylem fibers than in xylem vessels (Kim and Daniel, 2012). This higher intensity of mannans in the xylem fiber cell wall could suggest that these compounds are more important to mechanical support than water conduction (Kim and Daniel, 2012).

### Disruption of SWEET11 and SWEET12 expression differentially affects the cell wall composition of different xylem cell types

To further understand how modifications in facilitated sugar transport influence xylem secondary cell wall formation, we acquired Raman spectra for different xylem cell types from the *sweet11-1sweet12-1* double mutant (Fig. 1). As previously described, CART-based classifications were built to compare the different xylem cell types from both genotypes. For each cell type (VV, VF or FF), the original dataset was split into a calibration dataset and a validation dataset, after which the CovSel algorithm was applied on the calibration dataset to select the 10 Raman shifts showing maximum covariance. The CART models were then built on the calibration datasets and later applied on the validation datasets (Table 4, 5 and 6). The resulting CART tree classifications are displayed in Fig. 5. Regarding the cell walls between adjacent xylem vessels, two Raman shifts, namely, 1001 and 1093 cm^-1^, were sufficient to differentiate wild-type spectra from the double mutant spectra (Fig. 5A). Interestingly, these two Raman shifts have been shown to be associated with cellulose compounds (Gierlinger and Schwanninger, 2007; Özparpucu *et al*., 2017). Model performance was estimated for both calibration and validation datasets (Table 4), with the results demonstrating that the model can accurately discriminate spectra from both genotypes since the overall accuracy, sensitivity, specificity, PPV and NPV calculated for the validation dataset were between 76% and 90 % (Table 4).

**Fig. 5.**
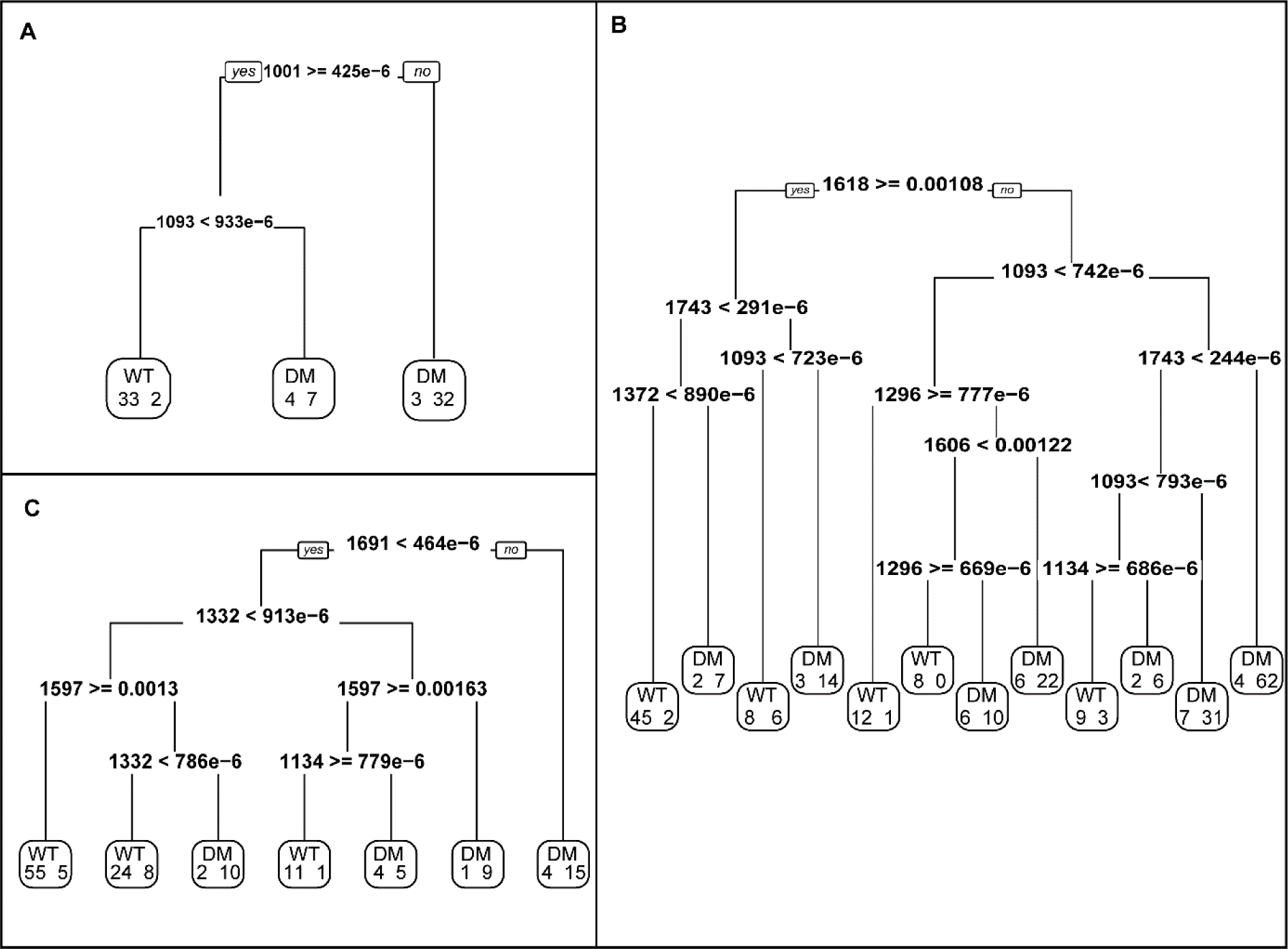
CART classification of wild-type and *sweet11-1sweet12-1* xylem Raman spectra. The classification trees have been generated by the CART method after ten-fold crossvalidation of the calibration dataset model, which was built using the 1800-1000 cm^-1^ range from the Raman spectra of cell walls between two xylem vessels (A), between a xylem vessel and a fiber (B) and between two xylem fibers (C). The decision-making process involves the evaluation of if-then rules of each node from top to bottom, which eventually reaches a terminal node with the designated class outcome (WT: wild-type and DM: *sweet11-1sweet12-1*). The numbers in each terminal subgroup stand for the number of either WT or DM spectra.

**Table 4.** Classification results for using Raman shifts to predict which genotype cell walls between xylem vessels represent, with the model calibration dataset (80% of total dataset) using a ten-fold cross-validation method, and the validation dataset (20% of total dataset) using a CART-based algorithm. NPV: negative predictive value, PPV: positive predictive value.

|  |  |  |  |  |  |
| --- | --- | --- | --- | --- | --- |
| Xylem vessel<br>/Xylem vessel |  |  | Raman prediction |  | Accuracy |
|  |  |  | wild-type | *sweet11sweet12* | 88.9% |
|  | Calibration<br>model (after<br>ten-fold<br>cross-validation) | wild-type | 33 | 2 | PPV<br>94.3% |
|  |  | *sweet11sweet12* | 7 | 39 | NPV<br>84.8% |
|  |  |  | Sensitivity<br>82.5% | Specificity<br>95.1% |  |
|  |  |  | Raman prediction |  | Accuracy |
|  |  |  | wild-type | *sweet11sweet12* | 84.2% |
|  | Validation<br>model | wild-type | 10 | 1 | PPV<br>90.9% |
|  |  | *sweet11sweet12* | 2 | 6 | NPV<br>75% |
|  |  |  | Sensitivity<br>83.3% | Specificity<br>85.7% |  |

**Table 5.** Classification results for using Raman shifts to predict which genotype cell walls between a xylem vessel and fiber represent, with the model calibration dataset (80% of total dataset) using a ten-fold cross-validation method, and the validation dataset (20% of total dataset) using a CART-based algorithm. NPV: negative predictive value, PPV: positive predictive value.

|  |  |  |  |  |  |
| --- | --- | --- | --- | --- | --- |
| Xylem vessel<br>/Xylem fiber |  |  | Raman prediction |  | Accuracy |
|  |  |  | wild-type | *sweet11sweet12* | 84.8% |
|  | Calibration<br>model (after<br>ten-fold<br>cross-validation) | wild-type | 82 | 12 | PPV<br>87.2% |
|  |  | *sweet11sweet12* | 30 | 152 | NPV<br>83.5% |
|  |  |  | Sensitivity<br>73.2% | Specificity<br>92.7% |  |
|  |  |  | Raman prediction |  | Accuracy |
|  |  |  | wild-type | *sweet11sweet12* | 88.4% |
|  | Validation<br>model | wild-type | 17 | 6 | PPV<br>73.9% |
|  |  | *sweet11sweet12* | 2 | 44 | NPV<br>95.6% |
|  |  |  | Sensitivity<br>89.5% | Specificity<br>88% |  |

**Table 6.** Classification results for using Raman shifts to predict which genotype cell walls between xylem fibers represent, with the model calibration dataset (80% of total dataset) using a ten-fold cross-validation method, and the validation dataset (20% of total dataset) using a CART-based algorithm. NPV: negative predictive value, PPV: positive predictive value.

|  |  |  |  |  |  |
| --- | --- | --- | --- | --- | --- |
| Xylem fiber<br>/Xylem fiber |  |  | Raman prediction |  | Accuracy |
|  |  |  | wild-type | *sweet11sweet12* | 83.8% |
|  | Calibration<br>model (after<br>ten-fold<br>cross-validation) | wild-type | 90 | 14 | PPV<br>86.5% |
|  |  | *sweet11sweet12* | 11 | 39 | NPV<br>78% |
|  |  |  | Sensitivity<br>89.1% | Specificity<br>73.6% |  |
|  |  |  | Raman prediction |  | Accuracy |
|  |  |  | wild-type | *sweet11sweet12* | 73.7% |
|  | Validation<br>model | wild-type | 14 | 3 | PPV<br>82.3% |
|  |  | *sweet11sweet12* | 7 | 14 | NPV<br>66.7% |
|  |  |  | Sensitivity<br>66.7% | Specificity<br>82.3% |  |

Additionally, data obtained from the CART model produced using Raman spectra acquired for the walls between xylem vessels and fibers (VF spectra) show that seven Raman shifts - at 1093, 1134, 1296, 1372, 1606, 1618 and 1743 cm^-1^ – can be used to discriminate the genotypes (Fig. 5B). The 1093 cm^-1^ shift appears three times and the 1296 and 1743 cm^-1^ shifts appear twice in the CART tree, suggesting that these three shifts are most important to discriminating the two genotypes (Fig. 5B). Based on the literature, the 1093 and 1372 cm^-1^ shifts are related to cellulose (Chylinska *et al*., 2014; Özparpucu *et al*., 2017), while the 1743 cm^-1^ is assigned to the υ(C=O) ester in pectins or hemicelluloses compounds (Chylinska *et al*., 2014). The 1606 and 1618 cm^-1^ Raman shifts have been reported to describe lignin bands (Prats Mateu *et al*., 2016). The performance of the model produced from VF spectra (for both calibration and validation datasets) was similar to the previous model.

Finally, the CART tree built using the spectra acquired for the walls between adjacent xylem fibers (FF spectra) shows that four Raman shifts can be used to discriminate between the wild-type and double mutant spectra. These shifts occur at 1134, 1332, 1597 and 1691 cm^-1^ (Fig. 5C), and are combined differently to distinguish both genotypes. The 1332 and 1597 cm^-1^ shifts are used twice in the classification trees, suggesting that both of these shifts are important to differentiating the WT xylem fiber cell wall from the DM xylem fiber cell wall (Fig. 5C). Interestingly, both bands are related to lignin compounds (1332 cm^-1^: aliphatic O-H bending; 1597 cm^-1^: aromatic ring stretching) (Özparpucu *et al*., 2017). The different descriptors for the models built from the calibration and validation datasets range from 67 to 89%, suggesting that the models can be used to distinguish the xylem fiber cell wall compositions of the two studied genotypes (Table 6).

To summarize, it seems that Raman shifts that discriminate cell walls between xylem vessels of the two genotypes are related to polysaccharides while the cell walls between xylem vessel/fibers and between adjacent fibers of the two genotypes can be discriminated using both lignin and polysaccharides shifts. The application of a CART-based analysis enabled us to identify new Raman shifts that could be used to better characterize the double mutant in a cell-specific manner. Interestingly, our previous analysis of the *sweet11-1sweet12-1* double mutant xylem cell wall did not reveal modifications in lignin composition (Le Hir *et al*., 2015). These discrepancies can be explained by the application of Raman microspectroscopy, which provided a spatial resolution of 2 μm x 2 μm, and therefore, the possibility to investigate individual cell types. The precision offered by Raman microspectroscopy suggests that the differences observed between wild-type and the *sweet11-1sweet12-1* mutant line could depend on the xylem cell type analyzed. Recently, Smith et al. (2017) demonstrated that xylem vessel lignification in the Arabidopsis floral stem is a non-cell-autonomous process that relies on the monolignol exchanges between xylem parenchyma or fiber cells and developing xylem vessels. If this model is extended to our research, it could be suggested that in addition to monolignols, sugar exchanges - mediated by SWEET facilitators - between xylem parenchyma cells and developing xylem vessels could also be commonplace in Arabidopsis.

## Conclusion

Synchrotron radiation FTIR and Raman spectroscopy are powerful tools for studying cell wall composition in plants both at the tissue and cellular level. Here, the application of SR-FTIR allowed us to picture the cell wall composition of phloem tissue in the Arabidopsis floral stem. Furthermore, CART-based classification, calculated using the Raman spectra acquired for different xylem cell types, identified spectral wavenumbers that could be leveraged to discriminate xylem cell types based also on their cellulose and hemicellulose composition. We also used both techniques to analyze the phenotype of the double mutant *sweet11-1sweet12-1*, which is deficient in the expression of two sugar facilitators that exist in vascular parenchyma cells. Our results showed changes in the hemicellulose composition of *sweet11-1sweet12-1* phloem cell walls when compared to WT plants. Moreover, analysis by Raman spectroscopy revealed that the disruption of both sugar transporters impacts xylem cell wall composition in a cell-specific manner. Therefore, SWEET11 and SWEET12 are important to ensuring correct phloem and xylem cell wall composition. Further addressing the role of SWEET facilitators in plant growth and development provides an attractive research direction that could provide answers for how intercellular sugar movements influence developmental processes such as vascular system development. Additionally, this study highlights that vascular parenchyma cells have a pivotal role in supplying the carbon skeleton required for cell wall formation in vascular tissues. The research approach presented here offers therefore the possibility of studying changes in cell wall polysaccharide composition at the cellular level and could be applied to investigations of how sugar transport affects cell wall formation in the vascular tissue of both herbaceous and ligneous species. Further, this approach open new perspectives for studying mutants affected in lignin biosynthesis and structure in the different xylem cell types, including, for instance, comparison between xylary fibers and interfascicular fibers.

## Supplementary data

**Fig. S1.** Illustration of the different spatial resolutions offered by vibrational spectroscopy techniques.

**Fig. S2.** Comparison of SR-FTIR peak areas of cellulose, hemicelluloses and lignin between xylem or phloem tissues from WT and *sweet11-1sweet12-1* plants.

**Fig. S3.** Average Raman spectra for the different xylem cell types in wild-type and *sweet11-1sweet12-1* lines.

## Acknowledgements

We thank Bruno Letarnec for his assistance in the greenhouse. The research was supported by the SOLEIL, the French national synchrotron facility (project n° 20150210). This work benefitted from a French State grant (LabEx Saclay Plant Sciences-SPS, ref. ANR-10-LABX-0040-SPS), managed by the French National Research Agency under an “Investments for the Future” program (ref. ANR-11-IDEX-0003-02). The authors declare no competing financial interests. The ChemFlow interface was supported by the Agropolis Foundation under the reference ID-1401-005 through the “Investissements d’Avenir” program (Labex Agro: ANR-10-LABX-0001-01).

